# Purification and Characterization of Inorganic Pyrophosphatase for *in vitro* RNA Transcription

**DOI:** 10.1101/2022.04.12.488088

**Authors:** Scott Tersteeg, Tyler Mrozowich, Amy Henrickson, Borries Demeler, Trushar R Patel

**Author notes:** These Authors contributed equally to this work (Scott Tersteeg, Tyler Mrozowich).

## Abstract

Inorganic pyrophosphatase (iPPase) is an enzyme that cleaves pyrophosphate into two phosphate molecules. This enzyme is an essential component of *in vitro* transcription (IVT) reactions for RNA preparation as it prevents pyrophosphate from precipitating with magnesium, ultimately increasing the rate of the IVT reaction. Large-scale RNA production is often required for biochemical and biophysical characterization studies of RNA, therefore requiring large amounts of IVT reagents. Commercially purchased iPPase is often the most expensive component of any IVT reaction. In this paper, we demonstrate that iPPase can be produced in large quantities and of high quality using a reasonably generic laboratory facility and that laboratory-purified iPPase is as effective as commercially available iPPase. Furthermore, using size-exclusion chromatography coupled with multi-angle light scattering and dynamic light scattering (SEC-MALS-DLS), analytical ultracentrifugation (AUC), and small-angle X-ray scattering (SAXS), we demonstrate that yeast iPPase can form tetramers and hexamers in solution as well as the enzymatically active dimer. Our work provides a robust protocol for labs involved with RNA *in vitro* transcription to efficiently produce active iPPase, significantly reducing the financial strain of large-scale RNA production.

**Statement of Significance:** We show an easy two-step purification procedure to efficiently produce large quantities of iPPase, an expensive component of IVT reactions. Laboratory-produced iPPase can significantly reduce the financial strain on research labs that rely on large-scale RNA production for experiments. Furthermore, we show for the first time, using a combination of orthogonal biophysical techniques, that yeast iPPase assembles into higher-order oligomers similar to bacterial iPPase.

## Introduction

Inorganic pyrophosphatase (iPPase; EC 3.6.1.1) is primarily characterized by its function to catalyze the hydrolysis of pyrophosphate into two monophosphates (1), as presented in equation 1. There are three different families of iPPase found in the cells: families 1 and 2 are free-floating in the cytosol, while family 3 or membrane-integral pyrophosphatases are membrane-bound in the mitochondria (2). The membrane-bound iPPase functions primarily as an ion pump, coupling the hydrolysis to move H^+^ or Na^+^ across the membrane. While membrane-bound iPPases are essential, family 1 and 2 iPPases are more abundant in cells. They are free-floating in the cytosol and help remove pyrophosphates from the cellular environment. Though families 1 and 2 perform similar functions, they are evolutionarily unrelated proteins (3). Family 2 has two domains per subunit, can hydrolyze approximately 2000 pyrophosphates per second, has a greater affinity to Mn^2+^ ions than other cofactors, and is predominantly found in bacterial species. Family 1, on the other hand, is found in all kingdoms of life, has a greater affinity to Mg^2+^ ions than other cofactors, can hydrolyze approximately 200 molecules per second, and only has one domain per subunit (4). Family 1 iPPase is the most studied iPPase and is the primary focus of our work.

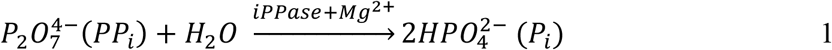

Family 1 iPPase is present in numerous organisms and displays variations in the structural arrangement. For example, the bacterial iPPase predominately exists as a hexamer, consisting of subunits of approximately 20 kDa (5). Yeast iPPase, however, forms a dimer in its active state with monomers of approximately 32 kDa (6). Family 1 iPPase is an essential enzyme for anabolic pathways such as RNA and DNA biosynthesis, and its removal has shown lethality in fungal cells (7).

IPPase is an essential component not only for cells to synthesize RNA but also for *in vitro* transcription (IVT) reactions. IVTs are expensive reactions that can produce large amounts of RNA, which is needed for almost all *in vitro* RNA biological experiments, including RNA structure-function characterization studies and the development of RNA-based therapies (8). IPPase drastically improves the yield of RNA through two different mechanisms (9). The first mechanism removes excess pyrophosphate produced during an IVT reaction, inhibiting IVT. The second mechanism prevents magnesium pyrophosphate formation (10). Magnesium ions are essential cofactors of T7 RNA polymerase, and without them, the functionality of T7 is dramatically reduced. By hydrolyzing the pyrophosphate before forming magnesium pyrophosphate, iPPase ensures that magnesium ions stay in the solution, optimizing T7 polymerase activity. With RNA therapeutics becoming increasingly popular, cost-effective methods must be developed to relieve the financial burden of IVT reactions so that more labs can perform research work on RNA-based systems.

This paper demonstrates that large-scale laboratory production of yeast iPPase is feasible, effective, and a viable alternative to commercially purchased iPPase. Additionally, by utilizing size-exclusion chromatography coupled with multi-angle light scattering and dynamic light scattering (SEC-MALS-DLS) and analytical ultracentrifugation (AUC), we show that yeast iPPase forms higher-order oligomeric tetramers and hexamers in solution as well as the commonly seen active dimer. Furthermore, we show the low-resolution structure of the iPPase tetramer in solution using small-angle X-ray scattering (SAXS) and the high-resolution structure via computationally predicted docking (SUPCOMB).

## Materials and Methods

### Overexpression and purification of yeast inorganic pyrophosphatase

Plasmid, pET29b-IPP1-His was a gift from Sebastian Maerkl & Takuya Ueda (Addgene plasmid # 124137). 15*µL* of Lemo21(DE3) E. *coli* cells were incubated with 10% volume of plasmid on ice for 30 minutes. The cells were then heat shocked for 45 seconds at 42*°C* after which 250*µL* LB media was added. The cells were incubated at 37*°C* for one hour and then plated on LB agar with Kanamycin (50 mg/mL) overnight.

Two 500 mL flasks of autoinduction media were induced with a single bacterial colony per flask and 0.1% volume of Kanamycin (50 mg/mL) and 0.1% of Chloramphenicol (50mg/mL). Bacterial cells were incubated with shaking at 37*°C* for 24 hours, followed by a 20*°C* incubation for 72 hours. All subsequent steps were performed at 4°C. The cell suspensions were harvested via centrifugation for 15 minutes at 5000 rcf. The total cell pellet (12.7 g) was resuspended in 30mL of lysis buffer (50 mM Tris, 100 mM NaCl, 5% glycerol, pH 8.0) along with 1 mM of PMSF, 5 mM of BME, 300 mg of lysozyme, and 0.1 mg DNase crystals (Thermofisher Scientific, Saint-Laurant, QC, Canada) and 250 mg of deoxycholic acid. Cells were lysed via sonication and then centrifuged at 30,000 rcf for 45 minutes. The supernatant was decanted and incubated with a 5 mL 50% slurry of NiNTA (Thermofisher Scientific, Saint-Laurant, QC, Canada) resin for one hour with constant inversion. The resin mixture was added to a gravity flow column, and the lysate was flowed through twice. 25 mL of wash buffer (50 mM Tris, 100 mM NaCl, 5% glycerol, 30 mM Imidazole, pH 8.0) was added to the column, and fractions were collected. Following the wash, 50 mL of elution buffer (50 mM Tris, 100 mM NaCl, 5% glycerol, and 500 mM Imidazole pH 8.0) was added to the column, with fractions collected similar to before. 10 *µL* of each collected fraction was mixed with 6X protein loading dye, heated to 95*°C* for 5 minutes, then loaded onto a 1.0 cm well PAGE casting plate (Bio-Rad Laboratories, Mississauga, ON, Canada). Finally, the samples were run on a 12% SDS-PAGE (sodium dodecyl sulfate poly-acrylamide gel electrophoresis) for 80 minutes at 150 V, followed by Coomassie staining, destaining, and visualization. Fractions with a pure protein band at the expected molecular weight were pooled and concentrated to 18 *µ*M using an Amicon Ultra-15 Centrifugal Filter Unit (10,000 Da MWCO) (Millipore Canada Ltd., Etobicoke, ON, Canada) in storage buffer (80mM Tris-HCl pH 7.5, 30mM MgCl_2_, 4mM spermidine, 20mM NaCl, 4mM DTT) and mixed 1:1 with 100% glycerol to a final concentration of 9 *µ*M and stored at -20°C in aliquots.

Affinity-purified iPPase was further purified using a Superdex 200 Increase 10/300 GL column (Global Life Science Solutions USA LLC, Marlborough, MA, USA) via an AKTA pure FPLC system (Global Life Science Solutions USA LLC, Marlborough, MA, USA) at 0.5mL/min, using iPPase storage buffer. Fractions were collected, and each peak was run on a 12% SDS-PAGE identically as described above.

### SEC-MALS-DLS

Light scattering experiments were performed on a Dawn® (Wyatt Technology Corporation, Santa Barbara, CA, USA) multi-angle light scattering instrument with 18 detector angles utilizing a 658 nm laser. Additionally, an Optilab® (Wyatt Technology Corporation, Santa Barbara, CA, USA) refractometer was fitted downstream to measure the solvent refractive index and provide the absolute concentration of iPPase. Upstream of both instruments was an SEC column (Superdex 200 increase 10/300 GL, Global Life Science Solutions, USA LLC, Marlborough, MA, USA) attached to an ÄKTA pure FPLC (SEC-MALS). Scattering experiments were performed at ambient temperature (25°C) with 0.5 mL/min flow rate and iPPase storage buffer (80 mM Tris-HCl pH 7.5, 30 mM MgCl_2_, 4 mM spermidine, 20 mM NaCl, 4 mM DTT). The refractive index of the solvent was defined as 1.042 (20°C), while the dn/dc (refractive index increment) value of 0.1850 mL/g was used for all iPPase measurements. A final concentration of 355 *µ*M iPPase was injected in a total volume of 500 µL. Data were processed and analyzed using Astra v8.0.0.25 (Wyatt Technology Corporation, Santa Barbara, CA, USA). Absolute molecular weight (Mw) was calculated using Equation 2, where: R(θ) is Rayleigh’s ratio, K is the polymer constant, and c is the concentration of the solute.

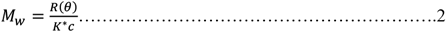

Integrated DLS measurements were taken every 3 seconds to determine the diffusion coefficient (Dτ) in the same scattering volume view by the MALS detector. To calculate the radius of hydration (R_H_), we used the Stokes-Einstein equation (Equation 3) (11). Whereas kB is the Boltzmann coefficient (1.380 × 10^−23^ kg·m^2^·s^−2^·K^−1^), T is the absolute temperature, and η is the viscosity of the medium (calculated to be 1.025). All values were calculated using ASTRA v8.0.0.25.

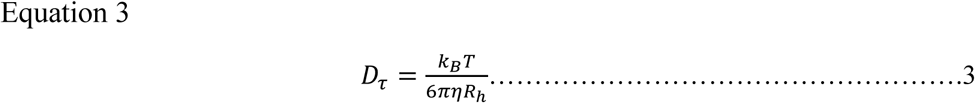

### Analytical Ultracentrifugation

AUC data for iPPase was collected via a Beckman Optima AUC centrifuge and an AN50-Ti rotor at 20°C. The iPPase sample was loaded at 0.5 OD at 280 nm (6.8 µM) and 0.5 OD at 220 nm (0.75 µM) into Epon-2 channel centerpieces in iPPase AUC buffer (80mM Tris-HCl pH 7.5, 30mM MgCl_2_, 20mM NaCl). Samples were centrifuged at 40,000 rpm, and scans were collected at 20-second intervals. We used the UltraScan-III package (11) to analyze sedimentation data via in-house supercomputer calculations. We analyzed the sedimentation velocity AUC data using two-dimensional spectrum analysis (2DSA) (12) with simultaneous removal of time-invariant noise, meniscus, and bottom positions fitted, as described previously (13), followed by enhanced van Holde-Weischet analysis (14). We estimated the buffer density and viscosity corrections with UltraScan (1.003g/cm3 and 1.0367 cP, respectively). Hydrodynamic parameters were corrected to standard conditions (20°C and water).

### Melting curve analysis

Melting curve analysis was performed on a Tycho (Nanotemper Technologies, Munich, Germany). Measurements were done in triplicate at multiple concentrations (50 µM, 5 µM, and 0.5 µM) in Tycho standard capillaries (Nanotemper Technologies). Data analysis was performed via MO Affinity Analysis Software v2.1.3 (Nanotemper Technologies, Munich, Germany).

### Small-angle X-ray scattering

Sample X-ray scattering data collection (HPLC-SAXS) was performed at the B21 beamline at Diamond Light Source (Didcot, UK), as described previously (15). Our iPPase sample (50 µL) was injected into an in-line Agilent 1200 (Agilent Technologies, Stockport, UK) HPLC connected to a specialized flow cell at 450 µM concentration and flowed through a buffer equilibrated Superdex 200 increase 3.2/300 GL (Global Life Science Solutions, USA LLC, Marlborough, MA, USA) size exclusion column at a flow rate of 0.160 mL per minute. Each frame was exposed to the X-rays for 3 seconds. Peak regions were buffer subtracted and merged via CHROMIXS (16). We analyzed the merged data using Guinier approximation to obtain the radius of gyration (R_g_) and study the homogeneity of samples (17). Dimensionless Kratky analysis (18) was also utilized to determine if the iPPase sample was folded, similar to previous examples (19). Furthermore, we utilized pair-distance distribution (P(r)) analysis via GNOM (20) to provide the R_g_ and the maximum particle dimension (D_max_). Using information derived from the P(r) plot, models were generated using DAMMIN (21), without enforced symmetry, as described previously (22). Finally, the previously generated models were averaged and filtered to obtain a single representative model through DAMAVER (23).

### High-Resolution Models with SAXS Envelopes

We performed CLUSPRO (24-29) docking analysis using our SAXS data as a constraint and PDB ID: 117e (30) (yeast inorganic pyrophosphatase) and generated 100 models. Using these 100 models, we then screened them against the raw scattering data using Crysol (31) to generate χ^2^ values based on the radius of gyration. Next, we visually inspected the top 5 results and overlaid them with our SAXS envelopes via Supcomb (32).

### In vitro transcription to evaluate iPPase activity

300 µL of 60 ng/µL Plasmid DNA was linearized prior to *in vitro* transcription via an Xba1 restriction digest site followed by heat inactivation (65°C for 20 minutes). IVT ingredients (Table 1) were mixed in the order listed to determine the functionality of iPPase produced in-house versus the iPPase that was previously purchased from Sigma Aldrich. T7 polymerase was produced and purified in-house, and Ribolock was purchased (Thermofisher Scientific, Saint-Laurant, QC). Samples were taken and quenched with an equal volume of 3M Sodium Acetate pH 5.3. After 4 hours at 37°C, the reaction was quenched with an equal volume of 3M Sodium Acetate pH 5.3. 10μL of each time point were mixed with RNA loading dye and heated to 95°C before loading on a 1.0 cm well PAGE casting plate (Bio-Rad Laboratories, Mississauga, ON, Canada). Samples were run on a 7.5% Urea PAGE gel for 30 minutes at 300V in 0.5X TBE buffer. The resulting gel was stained with SybrSafe (Thermofisher Scientific, Saint-Laurant, QC) and visualized.

**Table 1:** Components and final concentrations of the *in vitro* transcription reaction. *T7 Polymerase produced in-house.

| **<u>COMPONENT</u>** | **<u>FINAL CONCENTRATION</u>** |
| --- | --- |
| **5X TRAB** | 1X |
| **DTT** | 10mM |
| **NTPS** | 3mM |
| **GMP** | 5mM |
| **IPPASE** | 0.29µM (Purchased)/0.18µM (Lab) |
| **\*T7 POLYMERASE** | 0.18µM |
| **RIBOLOCK** | 0.5% (v/v) |
| **PLASMID DNA** | 18 ng/µL |
| **MILLIQ** | Up to desired volume |

## Results

### Purification of Yeast Inorganic pyrophosphatase

Yeast Inorganic pyrophosphatase was expressed in Lemo21(DE3) *E. coli* cells, followed by the initial purification using affinity chromatography (Figure 1A). The elution fractions show a high degree of expression of a band at the 35 kDa ladder size, which is consistent with monomeric iPPase (32 kDa). Elution 1 also contains a higher band at ∼66 kDa and some minor contaminating species. Next, to ensure that the iPPase was homogeneous for downstream studies, affinity-purified preparation was subjected to size-exclusion chromatography (SEC). The SEC chromatogram in Figure 1B shows one predominant peak eluting at ∼13.5 to 14.5 mL and two minor peaks eluting at ∼11-12 mL and ∼12-13 mL. These peak fractions were collected and further analyzed on an SDS-PAGE. As seen in Figure 1C, all three peaks contain a distinct species of approximately 35 kDa in size, similar to the sequence molecular weight of iPPase (32kDa) and consistent with affinity chromatography. Peak 3, similarly to affinity chromatography, shows an additional faint band above the monomeric iPPase species just above the 65 kDa ladder. Figure 1D shows the results of urea-PAGE analysis of in vitro transcribed RNA using commercially purchased iPPase (stock) and lab expressed and purified iPPase. Both T_F_ (time final) lanes show a similar amount of RNA being produced and a similar quality of RNA.

**Figure 1.**
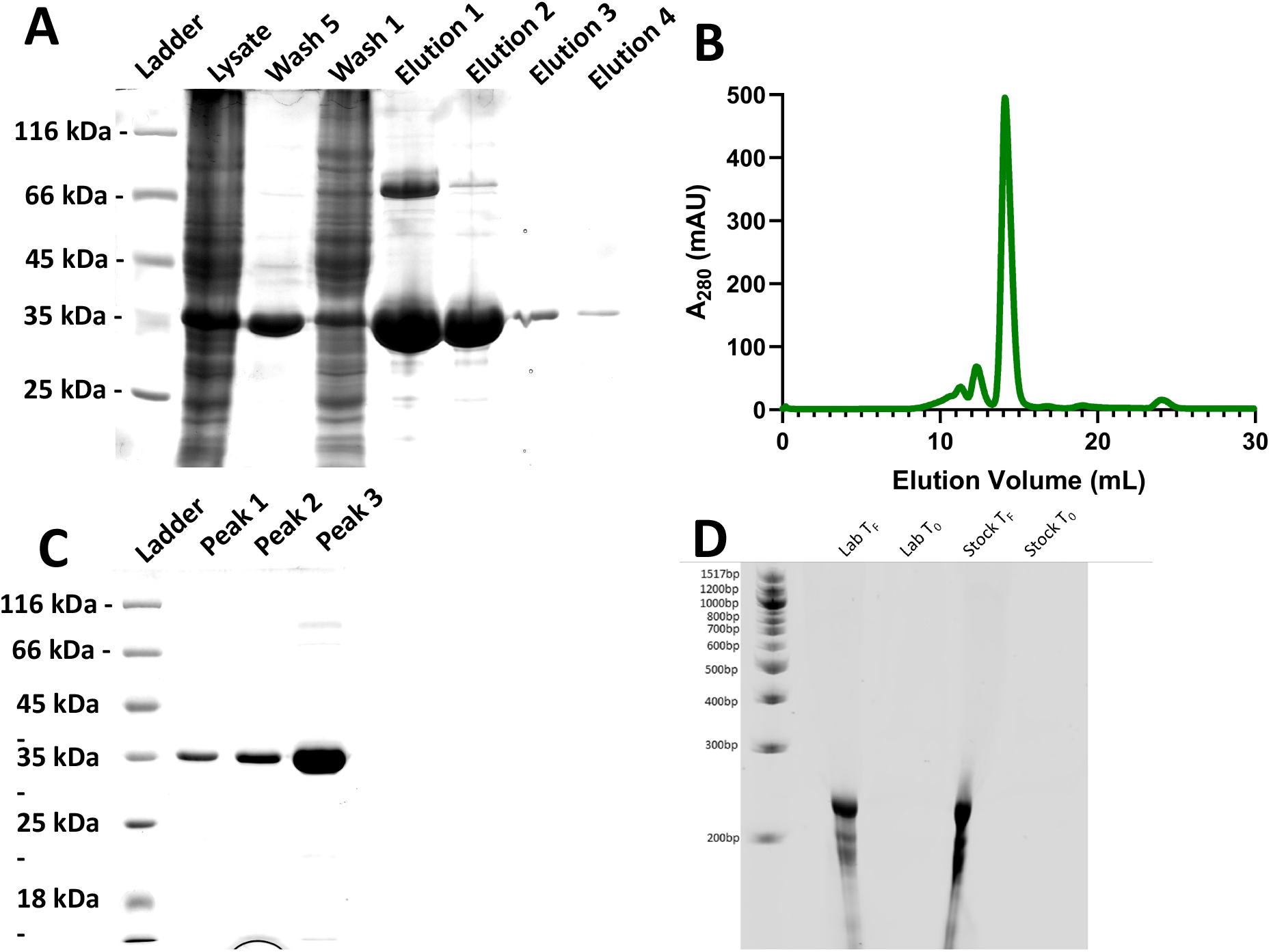
Purification of yeast iPPase. (A) SDS-PAGE after affinity purification via Ni-NTA. (B) Size-exclusion chromatogram purification of pooled elutions from after affinity purification. (C) SDS-PAGE after size exclusion chromatography. (D) RNA preparation using in vitro transcription demosntrates that the laboratory-purified iPPase is active. T_F_ (time final) lanes show a similar amount of RNA being produced with laboratory-purifired and the commerically purchased (stock) iPPase.

### Biophysical Characterization of Yeast iPPase

As we observed two additional peaks in Figure 1C, we were curious to investigate the possibility of iPPase forming oligomeric conformations in solution. Therefore, the affinity purified iPPase was subjected to SEC-MALS-DLS analysis. By coupling MALS and DLS with SEC, we obtained the absolute molecular weight for the three different peaks and the radius of hydration (R_H_), which are essential parameters for defining the size and shape of the proteins. The profile from SEC-MALS-DLS displayed similar trends as Figure 1B. As presented in Figure 2A, the peak eluting at ∼13.5 mL-15 mL has a molecular weight of 63.78 ± 1.54 kDa. Based on the sequence Mw of 32 kDa, this is consistent with yeast iPPase, which forms a functional dimer in solution. The peak eluting at ∼12 mL-13 mL has a molecular weight of 125.00 ± 3.20 kDa, which is the predicted weight of a tetramer. The final peak eluting from ∼11 mL-11.5 mL has a molecular weight of 186.90 ± 5.45 kDa, the weight of predicted hexameric iPPase. Figure 2B presents the R_H_ measurements derived from DLS with the peak eluting from ∼13.5 mL-15 mL, corresponding to dimeric iPPase, with an R_H_ of 3.56 ± 0.08 nm. The peak eluting at ∼12 mL-13 mL, corresponding to tetrameric iPPase assembly, has an R_H_ of 5.24 ± 0.14 nm. The hexamer peak eluted at ∼11 mL-11.5 mL, with an R_H_ of 6.31 ± 0.16 nm. As expected, the R_H_ values of iPPase increase as the size of the molecular assembly increases. Furthermore, the mass fraction % ratio of 83.3%, 11.6%, and 5.1% was calculated for the dimer, tetramer, and hexamer. Additionally, the mass fraction % ratio was converted to a molar ratio of the dimer, tetramer, and hexamer, resulting in a 55:3:1 ratio.

**Figure 2.**
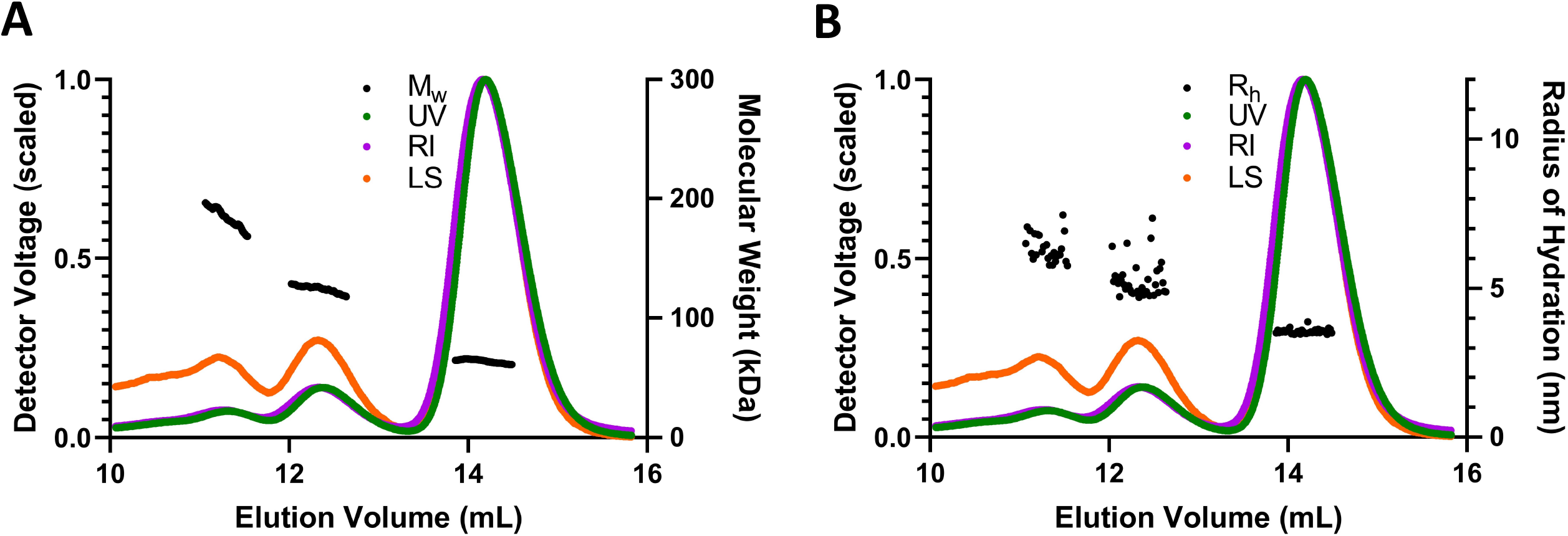
Light Scattering analysis of purified iPPase. (A) Absolute molecular weight analysis of purified iPPase via multi-angle light scattering. The black line(s) represent the molecular weight across each solute peak. (B) Hydrodynamic radius analysis of purified iPPase via dynamic light scattering. Black point(s) represent the hydrodynamic radius across each solute peak. R_H_ is the hydrodynamic radius, RI is the refractive index, UV is 280 nm, and LS is the light scattering signal.

The first orthogonal validation we chose was melting point analysis through A330/350 ratio measurements. With the Tycho NT.6, a protein can be quickly and easily analyzed by comparing the unfolding profile and structural integrity at different concentrations. Melting point analysis was performed on yeast iPPase at 0.5 µM, 5 µM, and 50 µM (Figure 3A). Each melting curve performed at different concentrations show the same pattern, with melting temperatures for 0.5, 5, and 50 µM of 64.0 ± 0.1, 63.9 ± 0.1, and 64.8 ± 0.1 °C, respectively. Additionally, we chose AUC as an orthogonal validation to confirm our experimental observations of oligomeric assembly. One of the advantages of using AUC is that biomolecules can be characterized at much lower concentrations than many other techniques such as SEC-MALS-DLS. As presented in Figure 3A, AUC analysis of affinity-purified (Figure 1B) carried out at 6.8 µM suggested the presence of three distinct peaks for iPPase in solution (Figure 3B). The sedimentation coefficient for the most prominent species is 4.5 S, followed by 6.5 S and 9.5 S for the tetramer and hexamer, respectively. Additionally, an enhanced VHW (van Holde-Weischet (14)) transformation was performed (Figure 3C) on two AUC runs at different concentrations (6.8 µM compared to 0.75 µM). These runs resulted in nearly identical VHW plots with most of the concentration around the 4.5 S value range, consistent with the dimer peak, and smaller concentrations extending above 6 S, consistent with the tetrameric and hexameric species. These results confirm that yeast iPPase can adopt quaternary arrangements in solution which differ from the enzymatically active dimer.

**Figure 3.**
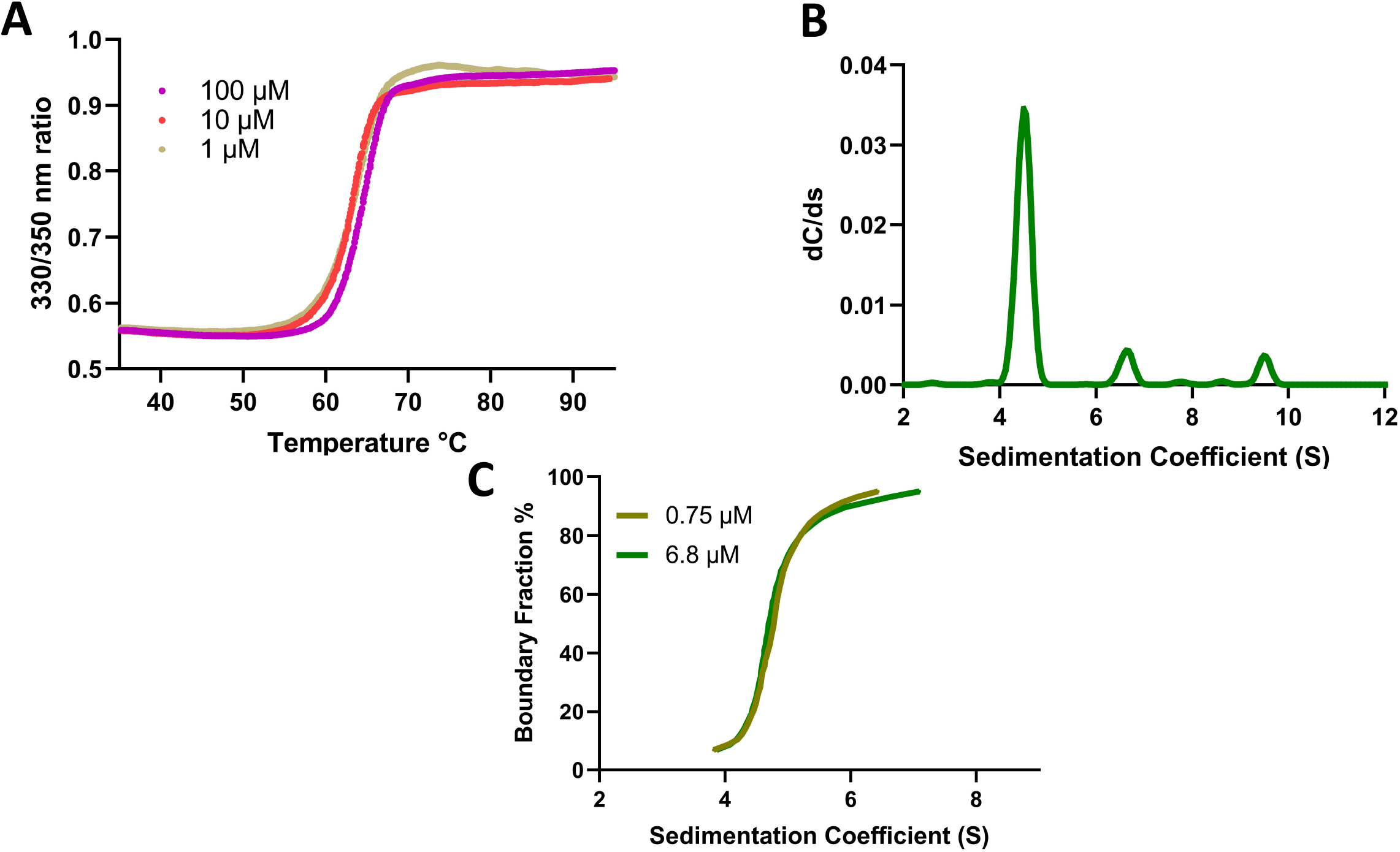
Melting curve and sedimentation-velocity analysis of iPPase. (A) Tycho NT.6 melting curve analysis of different concentrations of iPPase. (B) Sedimentation coefficient profile corresponding to iPPase dimer, tetramer, and hexamer. (C) Overlay of two enhanced van Holde-Weischet plots of 6.8 µM and 0.75 µM concentrations suggesting concentration independent oligomerization.

### Low-Resolution Structural Characterization of Yeast iPPase

Low-resolution structural information for iPPase was studied using affinity iPPase. First, the sample was injected into an HPLC instrument connected with the SAXS device to remove any aggregates. SAXS data from the prominent and tetrameric peaks was buffer subtracted, merged, and presented in Figure 4A. It should be noted that there was not enough homogenous scattering intensity to proceed forward with the processing of the hexameric peak. The merged data was processed using the Guinier method (plot of (I(q)) vs. (q^2^)), which allows for analysis of purity and determination of the R_g_ from the low-q region data (17). Figure 4B represents the Guinier plots for both the iPPase dimer and tetramer, with the low-q data linearity demonstrating that both populations are monodispersed and free of aggregation. Guinier analysis resulted in R_g_ values of 28.16 ± 0.04 Å and 45.77 ± 0.41 Å for iPPase dimer and tetramer, respectively (Table 2). Once the monodispersity of each peak was confirmed, the SAXS scattering data from Figure 4A was further processed to obtain dimensionless Kratky plots allowing for the detection of the relative foldedness of each population (33,34). The dimensionless Kratky plot for the dimeric state demonstrates it is well folded and globular (18) in solution with a maximal y-axis value of 1.15 ((I(q)/I(0)*(q*R_g_)^2^) (Figure 4C). The tetrameric state demonstrates a y-axis maxima value of 1.5.

**Figure 4.**
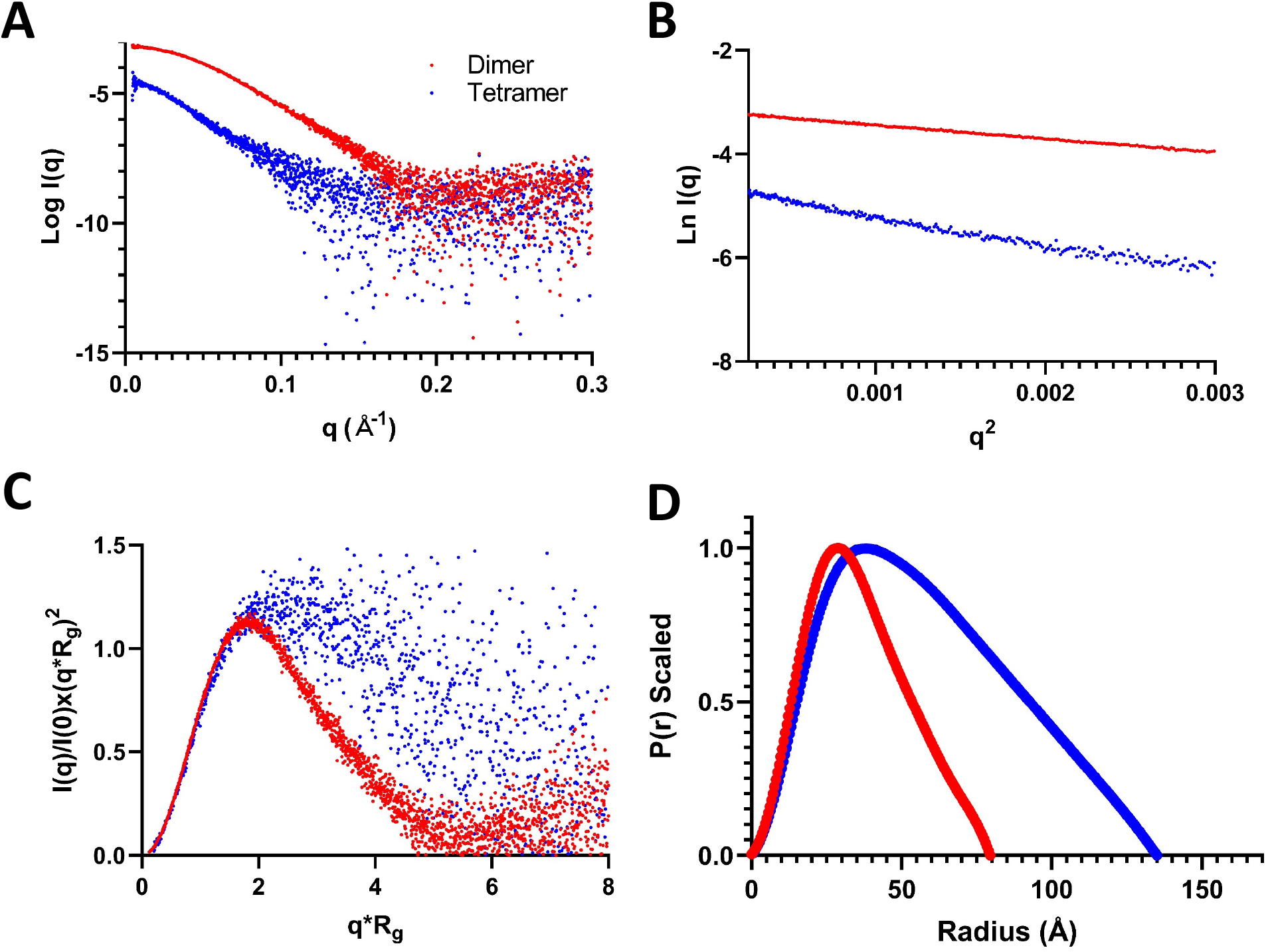
Characterization of iPPase oligomerization via SAXS. (A) Scattering intensity (Log *I*(*q*)) vs. scattering angle (q = 4πsinθ/λ) represents merged SAXS data. (B) Guinier analysis (*ln*(*I*(*q*)) versus *q*2) allows for homogeneity interpretation and determination of R_g_ via the low-angle region data. (C) Dimensionless Kratky plots (*I*(*q*)/*I*(*0*)*(*q*\**Rg*)2 vs. *q*\**Rg*) demonstrate that iPPase folds into a relatively globular state(s). (D) Pair-distance distribution (P(r)) plots for both iPPase peaks represent their maximal particle dimensions and allow R_g_ determination from the entire SAXS dataset.

**Table 2:**
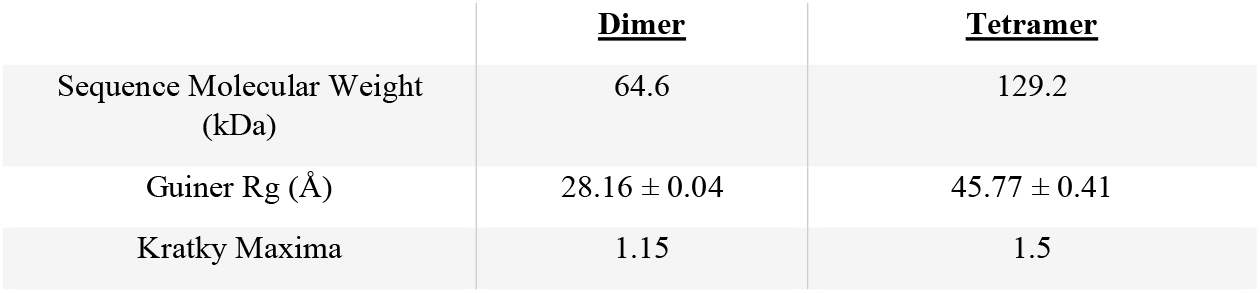
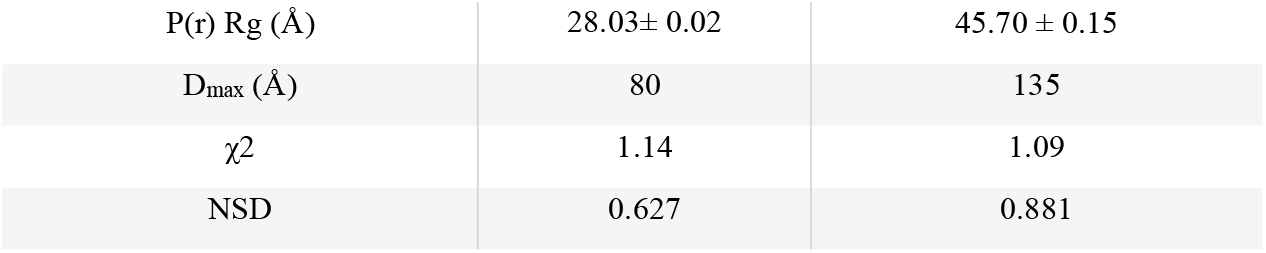
Small-angle X-Ray scattering parameters.

Next, the data was transformed via an indirect Fourier transformation to convert the reciprocal-space information of Figure 3A into the real space electron pair-distance distribution function (P(r), Figure 4D). Using the P(r) plot, R_g_ and D_max_ for both oligomeric states were calculated. P(r) analysis resulted in D_max_ values of 80 Å and 135 Å for the iPPase dimer and tetramer, respectively, and 28.03 ± 0.02 Å, 45.70 ± 0.15 Å, for the R_g_ values for the iPPase dimer and tetramer respectively. These R_g_ values are highly similar to the ones provided by the Guinier analysis, indicating an excellent fit of the data. Furthermore, the shape of the P(r) plot can indicate the solution conformation, with the iPPase dimer displaying an expected Gaussian shape, typical for globular proteins. The iPPase tetramer displays a right-side skewed Gaussian distribution, indicating an extended conformation.

DAMMIN was utilized to generate low-resolution 3-D structures of the iPPase dimer and tetrameric oligomerization states. We calculated 12 models for both the iPPase dimer and tetramer with a good agreement with the experimental scattering data and calculated scattering data. The χ2 values for both cases were ∼1.1, representing an agreement between the experimentally collected and low-resolution model-derived scattering data (Table 2). Next, DAMAVER was used to rotate and align all models, obtaining an averaged filtered structure for each oligomeric state (Figure 5). For each state, the goodness of the superimposition of individual models was estimated by the overlap function; normalized spatial discrepancy (NSD). The NSD value for the model agreement of the iPPase dimer was 0.627 and 0.881 for the tetramer, suggesting a good agreement between the models of each oligomeric state. Figure 5 presents both average filtered structures for the dimeric and tetrameric organization of iPPase and shows different orientations of each structure.

**Figure 5.**
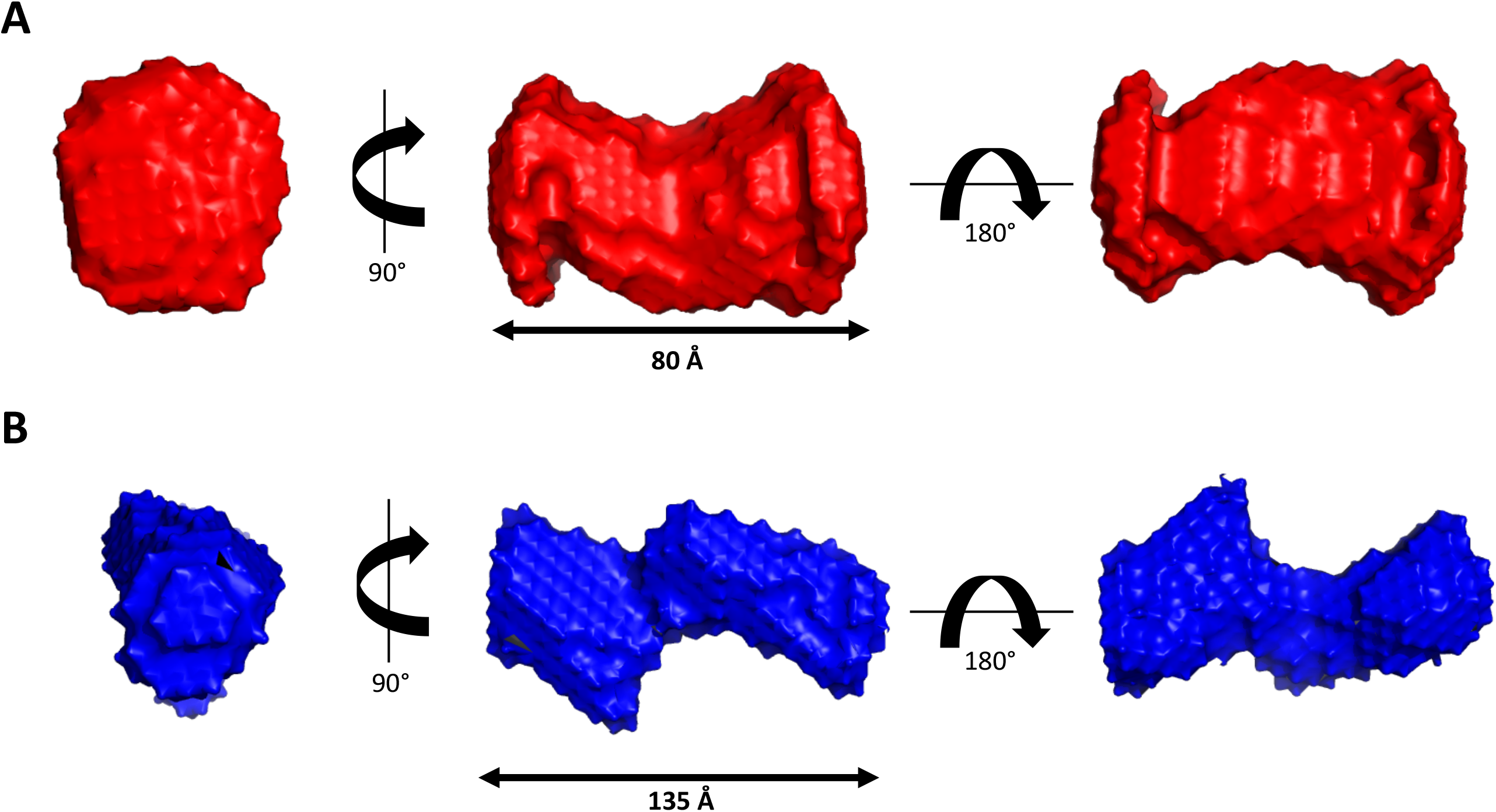
Low-resolution structural representations of iPPase oligomerization. (A) Three structures representing a 90° rotation about the y-axis and a 180° rotation about the x-axis from the middle representation of the iPPase dimer. (B) Three structures representing a 90° rotation about the y axis and a 180° rotation about the x-axis from the middle representation of the iPPase tetramer.

### High-Resolution Characterization of Yeast iPPase tetramerization

After calculating low-resolution structures for iPPase, calculations were performed to fit the known crystal structure into the generated low-resolution models. DAMSUP was utilized to orient the known crystal structure (PDB ID:117e) into our low-resolution structure. Visually, the high-resolution structure fits well with our low-res structure (Figure 6A). Next, CLUSPRO was utilized to dock the known iPPase dimer crystal structure into a tetrameric organization generating 100 possible models. We screened these 100 possible models against our raw scattering data, choosing the model with the lowest χ2 value (1.41), and overlaid it against our low-resolution model (Figure 6). Like the iPPase dimer, the tetramer also visually displays a good fit.

**Figure 6.**
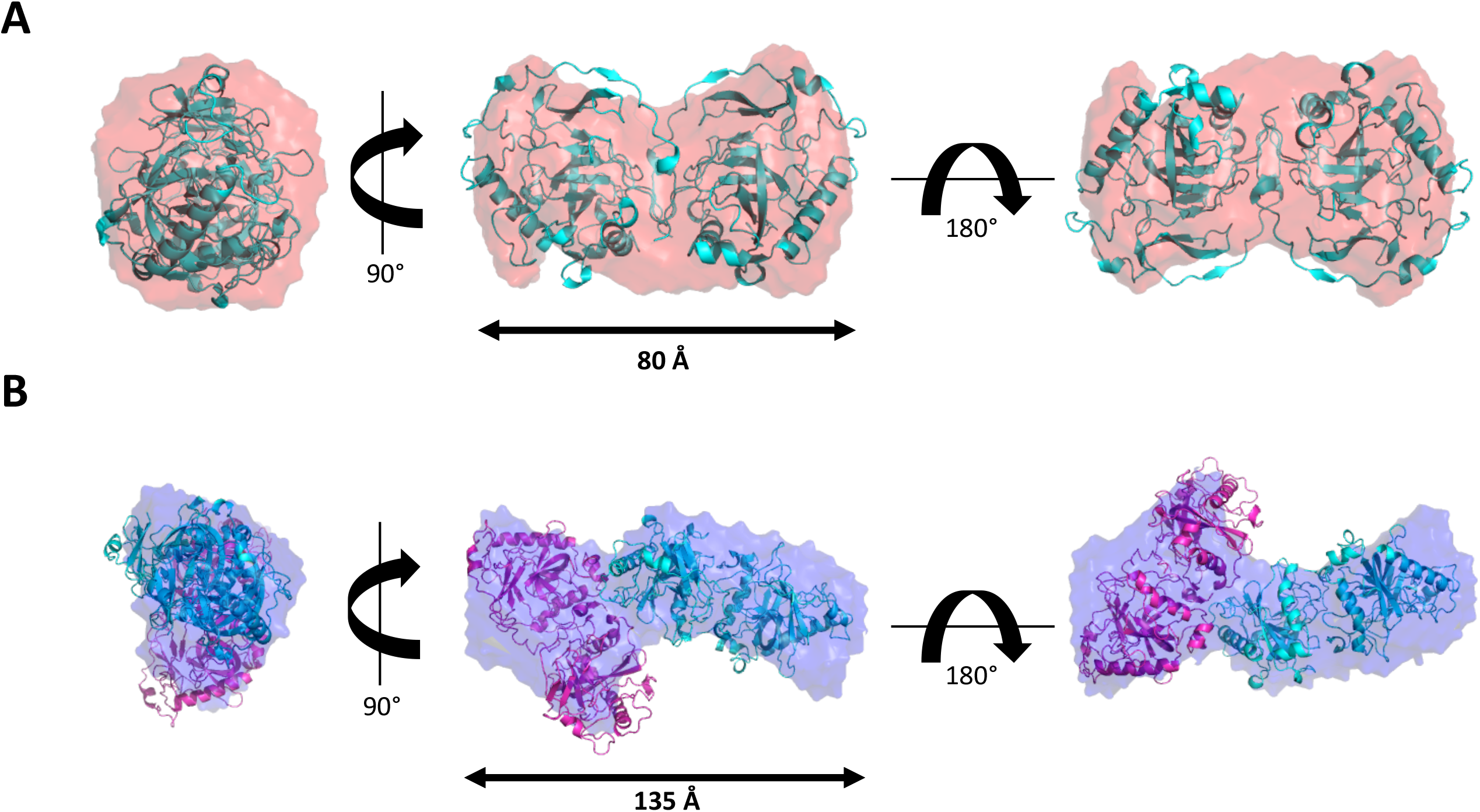
High-resolution representations of the iPPase dimer and tetramer. (A) Three structures representing a 90° rotation about the y axis and a 180° rotation about the x-axis from the middle representation of the iPPase dimer with an overlayed high-resolution crystal structure. (B) Three structures representing a 90° rotation about the y axis and a 180° rotation about the x-axis from the middle representation of the iPPase tetramer with an overlayed high-resolution crystal structure docked via ClusPro.

## Discussion

As iPPase is an essential enzyme for IVT reactions, it is paramount that any overexpression and purification will need to result in a large quantity of pure product. Initial affinity purification resulted in a large quantity iPPase (Figure 1A); however, further confirmation was needed with additional bands being present. The affinity-purified iPPase was subjected to a common additional purification step; size exclusion chromatography. The protein was processed through an S200 SEC column and visualized on an SDS-PAGE. Figures 1B shows three distinct peaks in the SEC chromatogram, and when visualized, all three peaks displayed the same size (figure 1C). Peak 3, which we believed to be the dimeric iPPase, still showed a larger protein species similar to affinity purification, which we believed to be a small amount of not fully denatured iPPase dimer.

A test IVT was performed to evaluate if the lab-purified iPPase is active and works well compared to the commercially available iPPase. As presented by the Urea-PAGE (Figure 1D), there was no difference in the amount and quality of RNA produced between the purchased and lab purified iPPase. Additionally, a lower concentration of lab-made iPPase 0.18 µM was utilized than the commercially purchased iPPase (0.29 µM). This concentration is promising for labs involved with RNA preparation as overall, the lab-made iPPase is just as effective as commercial vendors but a lot cheaper to prepare and purify.

The ambiguity of this additional protein band led us to further biophysical characterization through AUC, SEC-MALS-DLS, and eventually SAXS. First, we chose a combination of MALS and DLS coupled with size exclusion chromatography. MALS will importantly yield the absolute molecular weight across a peak when coupled to SEC, and DLS will gather the radius of hydration from the same peak. Therefore, utilizing SEC-MALS-DLS, the most prominent peak had a molecular weight that matched the dimer of iPPase (63.78 kDa) (Figure 2A). This result was expected, as it is known that yeast iPPase is active as a dimer (6). The middle peak gave an absolute molecular weight very close to a tetramer of iPPase (125 kDa). This was a surprising result because iPPase has not been characterized to exist in tetrameric conformation. Finally, the first eluted peak has a molecular weight very close to what would be the hexamer (186 kDa). This observation is interesting because it has been shown that bacterial iPPase forms an active hexamer (5), but it has never been shown that yeast iPPase can form any other tertiary structures apart from dimers. Next, the SEC-DLS experimental results were analyzed. These results corroborated the SEC-MALS results showing that while each SEC peak showed a single band on SDS-PAGE, corresponding to the monomer molecular weight, they have different hydrodynamic radii (Figure 2B). The third peak corresponding to the dimeric form had an R_H_ of 3.6 nm, consistent with proteins of similar size such as BSA (35). The second and third peaks, corresponding to the tetramer and hexamer, resulted in R_H_ of 5.24 and 6.31 nm, respectively, increasing as molar mass increased. Therefore, based on SEC-MALS-DLS, there is strong evidence that yeast iPPase forms both tetramers and hexamers in solution and the active dimer. However, with the large concentration used in SEC-MALS-DLS, these observations might be artificially induced, so an orthogonal approach was needed, which could be performed at a lower concentration.

The first orthogonal validation we chose was melting point analysis via Tycho NT.6. With the Tycho NT.6, a protein can be quickly and easily analyzed by comparing the unfolding profile and structural integrity at different concentrations (36). The Tycho NT.6 does this by comparing the ratio of intrinsic fluorescence at 350 nm and 330 nm, contributed by tryptophans and tyrosines. Therefore, if iPPase oligomerization was induced by high concentration, there should be a difference in the melting curves between different concentrations. Figure 3A presents the results at three distinct concentrations: 0.5 µM, 5 µM, and 50 µM. The melting curves for 5 µM and 50 µM present a very similar melting temperature of 64.0 and 63.9 °C, respectively, while 1 µM concentration resulted in 64.8 °C, which is still similar. Therefore, based on the melting temperature evidence, it does not seem that increasing the concentration of iPPase results in oligomerization.

To further verify that iPPase oligomerization is concentration-independent, AUC was chosen. AUC is a powerful technique, whereas biomolecules are subjected to extremely high centrifugal force and separated based on size, anisotropy, and density (37). AUC requires a relatively low concentration of the target sample (0.2-0.8 OD) and can be measured off-peak to further lower the target molecular concentration making it an ideal orthogonal method to pair with SEC-MALS-DLS. AUC analysis was performed at 6.8 µM, considerably less than the SEC-MALS analysis, and displayed the presence of three different quaternary structures in the pure iPPase sample. We can infer that the most prominent peak is the dimer as it has the highest concentration and the smallest sedimentation coefficient of 4.5 S. Then, as the species get larger, from tetramer to hexamer, the sedimentation coefficient increases from 6.5 to 9.5 S. To determine if the oligomerization was concentration-dependent, AUC experiments were performed at 220 nm, which allowed for much lower concentrations compared to 280 nm (0.75 µM vs. 6.8 µM). Figure 3C presents enhanced van Holde-Weischet plots showing boundary fraction % as a function of sedimentation. If iPPase oligomerization were concentration-dependent, the relative fraction of each state (dimer:tetramer:hexamer) would shift in response, and the plots would look significantly different. Both plots are congruent when overlaid, suggesting that oligomerization is not concentration dependent within the concentration range tested. With melting curve analysis, SEC-MALS-DLS, and AUC, this evidence suggests that the formation of different iPPase oligomeric structures is not dependent on concentration within the extensive range tested.

Next, we utilized SEC-SAXS to determine the low-resolution structure of iPPase in solution for multiple reasons. Firstly, a structural method was needed that could simultaneously separate biological molecules by size and allow the determination of structural features. Secondly, we needed a method in which an upper concentration limit would not be problematic. Concentration consideration was vital because it was determined that the molar ratio between the oligomeric states (dimer: tetramer: hexamer) is 55:3:1. Therefore, the amount of purified iPPase sample required to generate SAXS data would need to be enormous to ensure that the tetramer and the hexamer data would be usable. Unfortunately, while our injection of 450 µM was enough to get a large signal-to-noise ratio for dimeric and tetrameric peaks, we did not obtain a sufficiently high signal-to-noise ratio for the hexametric peak. This result is most likely due to the amount of protein loaded (13.28 mg/mL) above the SEC column recommended 10 mg/mL, which would broaden any peaks. Ultimately, we continued evaluating the dimer and tetrameric peaks, even though a usable hexameric peak was unattainable.

Initial SAXS analysis shows the difference in the concentration between the tetrameric and dimeric states by comparing how much tighter the data groups together as q increases in the dimer vs. the tetramer (Figure 4A). The Guiner plots also mirror this notion, with the low-q region of the dimer Guiner analysis showing virtually no data spread compared to the tetramer. Additionally, the Guiner analysis resulted in what we expected; the tetrameric oligomerization has a considerable increase in R_g_ (28.16 vs. 45.77 Å) (Figure 4B). This increase suggests that the oligomeric interface between the two stable dimers is likely an end-to-end interaction. This observation is further reinforced by comparing the dimensionless Kratky plots of both oligomeric states. The dimeric dimensionless Kratky has a y-maxima of ∼1.1, typical of globular proteins in solution. Comparatively, the tetramer has a dimensionless Kratky y-maxima of ∼1.5 and far less of a gaussian-like distribution (Figure 4C). This distribution is evidence of a more extended biomolecule, similar to nucleic acids (19,38-40). Comparing the D_max_ values for the dimer and tetramer (80 vs. 135 Å) reveals that the oligomeric interface is not perfectly end-to-end; if this were the case, the D_max_ would double (Figure 4D). However, since the D_max_ increases considerably, the oligomeric interface is likely an end-to-end interaction, not lengthwise. From the paired-distance distribution analysis of both oligomeric states, we observed that the real-space R_g_ values match almost identically with the reciprocal space R_g_ values derived from Guiner analysis for both the dimer and tetramer, respectively (28.16 vs. 28.93 Å and 45.77 vs. 45.70 Å). The similarity in R_g_ values suggests that both data sets are in excellent agreement and worthy of proceeding to 3-dimensional modeling.

3-Dimensional *ab inito* bead modeling was performed via DAMMIN, resulting in low-resolution structures for both dimeric and tetrameric iPPase. Figure 5 represents both the dimer and tetramer of iPPase in different orientations, showing that, as the previous data suggested, the tetramer seems to consist of two dimers in an extended conformation. The representative structures for dimeric and tetrameric iPPase oligomeric states studied have a χ^2^ value of ∼1.1, suggesting that the models have a good fit to the raw scattering data. Both dimer and tetramer are representative of a filtered average of 12 models with an NSD of 0.627 and 0.881, respectively. These NSD values show that the filtered models all have a very good fit to each other, and the representative model shown is an accurate representation of the model in solution.

Furthermore, since the high-resolution structure of yeast iPPase has been previously determined (30,41,42), we compared the high-resolution structure with our low-resolution SAXS models. As presented in Figure 6A, the high-resolution structure fits well into our solution scattering envelope, clearly showing both monomeric units of the crystalized homodimer. Overlaying high-resolution information into our SAXS envelope was an essential final step in validating our previous biophysical SEC-MALS and DLS data. Subsequently, we utilized CLUSPRO to dock the dimeric high-resolution crystal structure of iPPase to obtain 100 possible conformations of the iPPase tetramer. The docked models were screened using SAXS data and CRYSOL to determine which model is most likely the best representation of the tetrameric oligomerization state. We chose the model for a tetramer that agreed the most with the SAXS data, as evaluated by the lowest χ^2^ value, and represented it in Figure 6B. The high-resolution docked model fits very well visually with our SAXS envelope, validating our previous biophysical characterization that iPPase makes higher-order oligomeric species in addition to its biologically active homodimer.

Finally, we calculated the total cost of IVT reactions using a commercially purchased kit. One commonly used kit is the Ampliscribe™ T7-Flash™ Transcription Kit (Lucigen, Middleton, WI, USA) which sells for on average USD 241.00 for 25 reactions. If we scale this up to 1 mL IVT reaction, the approximate cost would be USD 482.00. Alternatively, when reagents are purchased separately, which include ATP, UTP, GTP, CTP (Sigma Aldrich, Oakville, ON, Canada), GMP (Sigma Aldrich, Oakville, ON, Canada), RiboLock (Thermo Fisher Scientific, High River, AB, Canada), in combination with in-house T7 polymerase and affinity-purified iPPase, the cost for 1 mL IVT reaction would be approximately USD 30.00. Thus, the laboratory-produced iPPase provides a cost-effective solution for large-scale IVTs. Finally, since it was determined that all three contaminant peaks in affinity-purified iPPase were oligomeric species of iPPase and that affinity-purified iPPase produced similar *in vitro* transcribed RNA, we believe that affinity purification alone is sufficient when using iPPase for in vitro transcription reactions.

Ultimately, this work provides a multi-faceted biophysical approach to characterizing the multiple oligomeric states of yeast iPPase. Furthermore, the expression and purification pipeline can provide a large quantity of active yeast iPPase to help offset high research costs incurred during RNA research. Ideally, a reduction in RNA research costs can significantly affect the ability of research labs to produce usable quantities of RNA and therefore reduce the barriers to doing RNA research.

## Author Contributions

Project was conceptualized by ST, TM, and TRP. ST and TM performed all experiments along with initial drafts of manuscript. AH and BD assisted in AUC experimental setup and analysis. All authors contributed to manuscript editing

## Acknowledgments

TM is supported by NSERC PGS-D fellowship. AH is supported by NSERC CGS. T.R.P thanks NSERC RTI and Canada Foundation for Innovation programs for infrastructure support. We thank DIAMOND Light Source B21 beamline staff for their help with data collection (SM26855). T.R.P is Canada Research Chair in RNA and Protein Biophysics, and B.D is Canada 150 Chair in Biophysics.

